# Warming and predation drive rapid evolution of ecosystem functioning but not functional traits

**DOI:** 10.64898/2026.03.17.712387

**Authors:** Laure Olazcuaga, Emma Couranjou, Laura Fargeot, Allan Raffard, Romain Bertrand, Murielle Richard, Kéoni Saint-Pé, Jérôme G. Prunier, Simon Blanchet

**Author notes:** These authors contributed equally to this work (Laure Olazcuaga, Emma Couranjou).

## Abstract

Global environmental change can rapidly reshape phenotypic trait distributions through adaptive and neutral processes. Yet, disentangling their relative contributions remains a major challenge, particularly for traits underpinning ecosystem functioning. Using a two-year mesocosm study, we experimentally evolved *Asellus aquaticus* under contrasting temperature and predation regimes to test how the joint influence of these global change pressures shape the evolution of traits and ecosystem functioning. We show that classic functional traits (body size and metabolic rate) showed no evidence of divergent evolution across evolutionary conditions, while ecosystem function itself, namely decomposition rate, has evolved rapidly and differently depending on evolutionary conditions, with the highest decomposition rate observed for *Asellus* populations exposed to high temperatures in the absence of their predator. A Pst/Fst comparison revealed that divergence in metabolic rate among evolutionary conditions was mainly driven by genetic drift, whereas, for body mass and decomposition rate, non-neutral processes (natural selection and/or plasticity) were also involved, albeit in a different way. By demonstrating that predation and temperature can influence the evolution of an ecosystem function over a few generations, this study represents an important first step toward uncovering the combined effects of global change on the eco-evolutionary dynamics of ecosystems.

## Introduction

Phenotypic traits are linking evolutionary and ecological dynamics, and understanding their spatial and temporal distribution in natural populations is a central endeavor. The distribution of phenotypic traits in natural populations is shaped by both deterministic and stochastic processes, including for instance selection, phenotypic plasticity, dispersal and/or genetic drift (Smith & Bernatchez, 2008). In turn, phenotypic traits variation (such as the distribution of individual body mass in populations) have consequences for ecological dynamics at the population and ecosystem levels as they determine when, where and how individuals use and transform resources (Woodward et al., 2005). Understanding the evolutionary processes shaping phenotypic trait distributions is crucial, as trait variability governs species responses to anthropogenic environmental changes (Palumbi, 2001). However, revealing these processes is challenging since deterministic and stochastic processes often act simultaneously in the wild.

Temporal and spatial environmental variations shape the evolutionary trajectories of populations. Populations can adapt to changing environments through selection acting on heritable phenotypic variability. This rapid adaptation based on genetic variation theoretically allows population size to maintain despite novel environmental conditions (Carlson et al., 2014; Gomulkiewicz & Holt, 1995). However, when environmental changes are strong and abrupt (i.e., when the selection pressure is strong), populations size can abruptly decline, which favors stochastic processes (genetic drift) and limit the adaptive potential of populations to the novel environmental conditions (Auld & Relyea, 2009; Orr & Unckless, 2008; Tanaka, 2000; Willi et al., 2006). In that case, the distribution of phenotypic traits in populations is expected to change randomly over time and space, which can lead to substantial phenotypic divergences among populations (Shama et al., 2011). Deterministic and stochastic processes often act simultaneously, producing a blend of adaptive and neutral phenotypic changes that is difficult to tease apart in natural populations (Hedrick & Garcia-Dorado, 2016; G. S. Stewart et al., 2017). For instance, predation by non-native species simultaneously leads to a reduction in population size of naïve prey and a selection against the most vulnerable phenotypes (Kitano et al., 2008). This type of change can be called a “selective bottleneck” because it involves both random changes due to reduced population size by predation, as well as adaptive changes due to selection imposed by the presence of the predator (Bassitta et al., 2021; Hulthén et al., 2024; Reger et al., 2018). However, when predation causes an excessive bottleneck, stochastic processes blur deterministic processes, which may ultimately lead to the extinction of the prey population (Patankar et al., 2006). The balance between stochastic and deterministic processes will depend notably on the type of selection pressure and its intensity (i.e., its consequence on the population size).

The change in phenotype distribution in a population following environmental change has often been studied (Violle et al., 2007). For instance, environmental change such as the introduction of new predation pressures has been shown to influence prey traits related to fecundity, such as body coloration and female mating preferences (Bierbach et al., 2011; Fisk et al., 2007). Some of these traits are thought to be tightly associated with ecosystem functions, such as primary productivity or litter decomposition, and are therefore referred to as functional traits (Hadj-Hammou et al., 2021; Portela et al., 2023; Violle et al., 2007). These functional traits include -amongst others- the body size and the metabolic rate of organisms, both of them being altered by global change pressures (Brown et al., 2004; Woodward et al., 2005). For instance, warming strongly affects individual metabolic rate as well as the distribution of body size in populations, especially in ectotherms (Daufresne et al., 2009; Stanislawek et al., 2025; Yvon-Durocher et al., 2010). However, few studies have focused on the evolution of ecosystem functions *per se*, which currently limit our ability to predict the eco-evolutionary consequences of global change on ecosystem functioning. Indeed, determining whether an ecosystem function and their underlying functional traits share similar evolutionary dynamics is key to predict the effect of new selective pressures on ecological dynamics (Govaert et al., 2021; Song et al., 2023; Wu et al., 2023).

Bridging ecology and evolution makes it imperative to investigate how functional traits and ecosystem function evolve (Streit & Bellwood, 2023; Violle et al., 2007). Yet, a major challenge remains to disentangle the relative roles of deterministic and stochastic processes, which often act simultaneously. Methods from quantitative genetics exist to independently measure the effects of drift and selection. Stochastic evolutionary mechanisms can be estimated by quantifying the level of neutral genetic differentiation (e.g., F_st_) among populations, using neutral genetic markers. An index of differentiation analogous to the F_st_ can be estimated from phenotypic traits (P_st_), which captures the degree of phenotypic divergences stemming from both stochastic and deterministic (selection and plasticity) processes. A direct comparison of the two indices (F_st_ and P_st_) provides a valuable and simple way for revealing the relative role of evolutionary processes on the distribution of phenotypic traits; when P_st_ is equal to F_st_, drift predominates, whereas when P_st_ is higher than F_st_, deterministic processes predominate (Merilä and Crnokrak 2001, Leinonen et al. 2008). These methods can further be applied to experimental evolution approaches, where environmental pressures are experimentally controlled, enabling causal links to be revealed (Kawecki et al., 2012). In particular, conducting this type of experimental approach in mesocosms allows approaching conditions close to those found in the wild, which is often a limit of fully controlled experimental evolution approaches (Pantel et al., 2015; R. Stewart et al., 2013; Xie et al., 2024).

Here, we aim to investigate how adaptive and neutral evolutionary changes (due to deterministic and stochastic processes respectively) under combined environmental changes affect the distribution of phenotypic traits and ecosystem functioning in a freshwater crustacean. More specifically, we address the following questions: (1) How are functional traits that supposedly support the ecosystem function influenced by the evolutionary conditions in which populations have evolved? (2) How is the actual ecosystem function influenced by evolutionary conditions? and (3) How do adaptive and neutral changes stemming from evolutionary changes influence these responses? To answer these questions, we measured traits in populations of *Asellus aquaticus* that have evolved for several generations under diverse environmental conditions in replicated mesocosms for two years. We manipulated two components of global change: (i) warming, by exposing crustaceans to +3°C conditions compared to the ambient climate (based on an intermediate climate change scenario, SSP2-4.5) and (ii) predator invasion by exposing (or not) crustaceans to a predatory fish *Phoxinus dragarum* (Miro and Ventura 2015). For all populations that evolved under these ‘evolutionary conditions’ (i.e., environmental conditions of the mesocosm in which the experimental populations evolved), we measured different traits. For functional traits, we focused on body mass and routine metabolic rate (measured across a temperature gradient to characterize thermal performance curves); for the ecosystem function, we focused on the ability of *Asellus* populations to decompose dead organic matter (also quantified along a thermal gradient). We expected an increase in (mass-corrected) metabolic rate and a decrease in body mass and size for experimental populations having been exposed to the warmer climate (Andreassen et al., 2025; Daufresne et al., 2009; Nespolo et al., 2011, respectively) or to predators (Auer et al., 2018; Reznick et al., 1990, respectively). We also expected that the ability to decompose organic matter will be higher for populations that have evolved under the warmest climate due to an increase in energetic needs (Martínez et al., 2014) and lower in populations that evolved under predator pressure due to behavioral adjustments (Raffard et al., 2017). Although phenotypic divergence among experimental *Asellus* populations is expected to be underlined simultaneously by both deterministic and stochastic evolutionary processes, we expected to observe an increase in the relative importance of stochastic processes in populations of *Asellus* having evolved under predation pressures compared to those that evolved under warm conditions, as predation may substantially reduce population sizes, hence further strengthening genetic drift (Bassitta et al., 2021; Reger et al., 2018). However, for both predation and warming, we expected selection pressure to lead to P_st_ values greater than F_st_, indicating that trait differentiation cannot be explained solely by genetic drift and suggesting the action of divergent selection between populations exposed and unexposed to predation.

## Materials and Methods

### 1. Model species

The benthic freshwater detritivore *Asellus aquaticus*, distributed throughout Europe and Asia, is an excellent model to investigate ecosystem dynamics and eco-evolutionary feedback between the environment and natural populations. This crustacean species displays a high potential for adaptation, supported by the large variation in phenotypes of populations living in contrasting environments (Hargeby et al., 2004; Lürig et al., 2019; Sworobowicz et al., 2015). *Asellus aquaticus* is particularly useful for tracking evolutionary changes because it has a relatively short generation time (three months to reach sexual maturity) enabling up to two complete generations per year (Lafuente et al., 2021). This species also has a good resistance to environmental variations, as evidenced by the broad distribution of the species (Aston & Milner, 1980; Maltby, 1995). Additionally, the position of *A. aquaticus* as a keystone species in freshwater ecosystems makes it a particularly interesting model to investigate the ecosystem dynamics. Indeed, *A. aquaticus* has a broad diet ranging from phytoplankton to leaf litter, and is predated by multiple species (e.g., fish, insects, and waterfowl; Hargeby et al., 2004; Hart & Gill, 1992; Lafuente et al., 2021). For the predation pressure experienced by our target species, we used the cyprinid fish *Phoxinus dragarum* (Occitan minnow). *P. dragarum* is a small-bodied cyprinid fish species having been widely introduced in lakes throughout Europe (Tiberti et al., 2022). It is a voracious fish species with a diverse diet that includes benthic macro-invertebrates such as *A. aquaticus* (Raffard et al., 2021). In addition, we included a warming treatment of +3 °C, as temperature is a key driver of physiological and ecological processes in aquatic ectotherms. The freshwater isopod *Asellus aquaticus* is sensitive to temperature variation, exhibiting optimal performance at relatively cool temperatures (∼15–20 °C) and reduced survival and performance at higher temperatures (>30 °C) (Bloor, 2010; Di Lascio et al., 2011).

### 2. Experimental evolution

#### a. Sampling and experimental evolution in mesocosm

Experimental populations of Asellus aquaticus have been created in mesocosm from the Aquatic Metatron (Richard et al., 2025) with individuals sampled in their natural habitat (Figure 1-A). Individuals were sampled in May 2019 from a unique population of origin located in a pond in Toulouse (South of France, 43°33’35.3‴N 1°28’26.6‴E). Each experimental population was created by introducing 50 individuals in each mesocosm tank. Mesocoms are 2m^3^ circular tanks filled with freshwater from the nearby Lez River and set to mimic a natural ecosystem. To do so, we set a gravel bed in each mesocosm and introduced various organisms (phytoplankton, zooplankton, invertebrates…) as detailed in Stanislawek et al. (2025). Mesocosms were distributed into four experimental evolutionary conditions, resulting from contrasting environments achieved by controlling two environmental pressures: ambient or warm (ambient +3°C) water temperature and presence or absence of the predator P. dragarum (Figure 1-B). The +3 °C warming treatment was chosen to represent an intermediate climate change scenario between moderate (SSP2-4.5, projecting a global temperature increase of ∼2.7 °C by 2100) and more severe climate projections (SSP3-7.0 and SSP5-8.5, projecting ∼3.6–4.4 °C by 2100).

**Figure 1.**
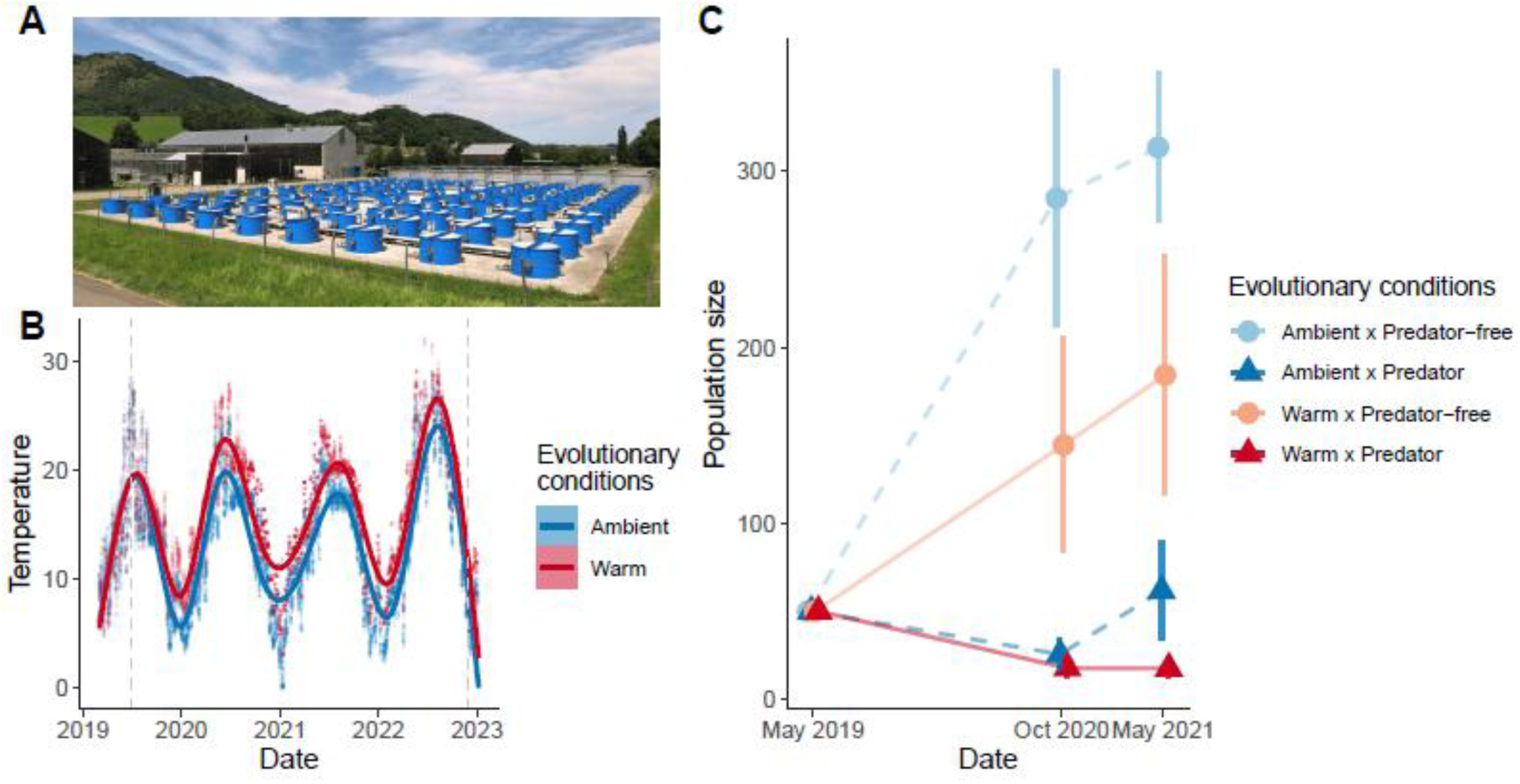
Experimental evolution in mesocosm. A) Picture of the Aquatic Metatron with the mesocosms used for this experimental evolution. B) Temperature dynamics throughout the experiment (x-axis: Year; y-axis: °C). C) Population size dynamics during the mesocosm experiment (x-axis: Month Year; y-axis: number of individuals). B) Each point shows the mean temperature recorded in an experimental tank. Lines represent generalized additive model (GAM) fits across all experimental populations evolving under warm (+3 °C; red) or ambient (blue) temperature conditions. C) Each point shows the mean population size per evolutionary conditions (± SE); colors correspond to the conditions in which populations evolved. The populations evolved either in warm temperature (red with a solid line) or ambient temperature (blue with a dotted line), either with the presence of a predator (dark colors with triangular points) or without a predator (predator-free; light colors with circular points). A slight jitter was added to avoid point overlap and enhance visibility.

From the 33 experimental populations created, 15 were used in this study, excluding those with insufficient population size (N_minimum_ = 10 individuals) while maintaining balance in the experimental design and into the 4 evolutionary conditions. Five experimental populations evolved under the ambient/predator condition, four populations under the warm/predator condition, four populations under the ambient/predator-free condition and two populations under the warm/predator-free condition. The conditions in which experimental populations have evolved will be hereafter referred to as “evolutionary conditions”, encompassing “temperature condition” and “predation regime”.

Each experimental population evolved from spring 2019 to winter 2022 (44 months, ∼6-8 generations, assuming two generations per year), under one of the four contrasting evolutionary conditions. Census data collected at two time points enabled us to track changes in population size (Figure 1-C). This monitoring confirmed that predator presence caused a marked decline in population size (dark lines compared to light lines in Figure 1-C).

#### b. Common garden

In December 2022, individuals from the experimental populations were sampled and transferred to a common garden prior to data collection. The purpose of this common garden was to minimize the effects of phenotypic plasticity due to evolutionary conditions. Prior to measurements, 290 individuals per experimental population were transferred into 300L aquaria at 20°C, for six weeks for metabolic rate and body mass and for two days for decomposition rate.

### 3. Data collection

To investigate how evolution of populations under contrasted evolutionary conditions impacts the distribution of traits, we measured the distribution of two functional traits: metabolic rate and body mass; and one ecosystem function: the ability of the population to decompose dead organic matter, measured as the leaf decomposition rate.

#### a. Metabolic rate

To assess the distribution of the metabolic rate under a temperature range of *A. aquaticus* in experimental populations, oxygen consumption was evaluated at 8 different test temperatures: 5°C, 10°C, 15°C, 20°C, 23°C, 25°C, 27°C and 30°C. For each temperature, the oxygen consumption of an average of 19 individuals per population was measured individually (a total of 152 individuals per experimental population:19 individuals per 8 test temperatures) by following the procedure described in Raffard et al. 2019. Briefly, each individual was placed into a 5mL PreSens vial SensorDishes® filled with water. Oxygen concentrations were collected continuously every 15 seconds for at least 2 hours with oxygen sensors within the vials. To account for acclimation of individuals, data from the first 20 minutes were removed before analyses.

A proxy for routine metabolic rate was obtained by measuring the slope of oxygen consumption over time, using linear regression (Svendsen, Bushnell, & Steffensen, 2016). A high coefficient of determination (r²) for this slope indicates high precision in the estimation of metabolic rate (Svendsen, Bushnell, Christensen, et al., 2016). Accordingly, only slopes with r² values exceeding a threshold of 0.85 were retained (McKenzie et al., 2000; Raffard et al., 2025). We confirmed that this strict threshold did not bias the results as analyses including all slopes produced consistent outcomes (Chabot et al., 2021). Consistent with standard practice (Chabot et al., 2016), we used the log-transformed absolute value of the slope coefficient of this relationship as a proxy for routine metabolic rate, hereafter referred to as *metabolic rate*.

#### b. Body mass and size

To estimate the individual body mass of *A. aquaticus* in experimental populations, 152 individuals from each experimental population were photographed individually to measure their body size and estimate their body mass. These 152 individuals corresponded to those whose metabolic rate was measured. Unlike metabolic rate, body mass was not estimated over a range of temperatures because momentary exposure to different temperatures is not expected to alter this trait. Body mass was estimated based on body size using a size-mass relationship. Indeed, body mass and body size in crustaceans typically follow an allometric power-law relationship (Maszczyk & Brzeziński, 2018; Peters & Wassenberg, 1983; Stevens, 2009), often modeled as body mass=a×body size^b^ after log–log transformation. Body size was measured for each individual by estimating the width of the base of the cephalothorax, just above the pereon with the ImageJ software (Schindelin et al., 2012). The size-mass relationship was obtained by measuring and weighting (dry mass of each individual to the nearest 10^-5^ gram) 57 individuals from all experimental populations (Figure S1). Then, the fitted allometric relationship (body mass = 765.99 × body size^3.08^) derived from the log–log model with an R² = 0.721 was used to infer the body mass of all photographed individuals. It is noteworthy that analyses presented below were done using both the measured body length and the estimated body mass, and they all provided the same conclusions. We are primarily interested here in body mass, but additional, similar analyses have also been carried out on body size.

#### c. Decomposition rate

To decipher how population evolution impacts the distribution of the decomposition rate across a temperature range, decomposition rate was assessed at 3 temperatures: 5°C, 15°C and 27°C by quantifying the mass of leaves degraded over time. For each test temperature and each experimental population, 3 groups of 10 individuals were placed in 3 water-filled plastic trays (300ml). In each plastic tray, we placed 1 gram of dry leaves (dried 48 hours at 60°C) of *Alnus glutinosa*, which is a common tree found in the natural habitat of *A. aquaticus*. After 17 days, the dry weight (dried 48 hours at 60°C) of leaf recovered in each plastic tray was measured. The decomposition rate was estimated as k = −(ln(X))/t (Raffard et al. 2021), where X is the proportion of litter remaining after and t the time elapsed in days. The different test temperatures (5°C, 15°C and 27°C), were kept constant by partially submerging trays in large aquariums (600L) whose water was either kept cold or warm using chillers and heaters respectively.

#### d. SNP sampling for F_ST_/P_ST_ analysis

Neutral genetic differentiation among populations (i.e., differentiation due to genetic drift) were estimated using a large panel of Single Nucleotide Polymorphism markers (SNPs) sampled randomly in the genome. DNA was extracted from eleven individuals in each experimental population using an extraction robot (externalized to Diversity Array, Australia). Thousands of SNPs were detected using a reduced-genome sequencing approach (DaRT sequencing, Jaccoud et al., 2001) and genotyped. SNPs processing (from library preparation to bioinformatics) were externalized to Diversity Array (Australia). This approach allowed the collection of 4425 polymorphic SNPs. DartSeq genomic data were then filtered by removing missing data and only keeping SNPs with less than 5% of missing data. To eliminate potential SNPs under selection, we used the OutFLANK package to assign a q-value to each locus to detect outliers that may be due to selection (Whitlock & Lotterhos, 2015). Based on this approach, no *F_ST_* outliers were detected, indicating that none of the SNPs were under selection (see Figure S2 for the relationship between *F_ST_* and heterozygosity). We therefore assumed that these SNPs followed a neutral dynamic (which is the case for most SNP panels developed using reduced-genome approaches, Martchenko & Shafer, 2023) and that this panel was adequate to estimate neutral differentiation.

### 4. Data analysis

All statistical analyses were performed in R version 3.4.2 (R Core Team, 2014). Linear mixed-effects models were fitted using the packages lme4 (Bates et al., 2015) and nlme (S. N. Pinheiro et al., 2020).

#### a. Trait variability: metabolic rate, body mass and decomposition

To test how body mass (or body size) was influenced by the evolutionary conditions in which experimental populations evolve, we fitted a linear mixed model with temperature conditions (two levels: warm and ambient) and predation regime (two levels: predator and predator-free) as fixed effects. To take the variation among replicated experimental populations into consideration, we included experimental population identity as a random effect.

To test how metabolic rate and decomposition rate were influenced by the evolutionary conditions in which experimental populations evolve and the environment in which individuals were measured (test temperature), we extended this model by adding the test temperature as a covariate (continuous variable). For decomposition rate, test temperature was included as a linear covariate. For metabolic rate, the relationship with test temperature was non-linear; we therefore modelled it using a natural cubic spline with three degrees of freedom, which allows for flexible curvature beyond a simple quadratic term.

For metabolic rate, we also integrated the log-transformed mass of individuals as covariates to correct oxygen slopes over time by the log of the mass of individuals and thus to refer to metabolic rate (Raffard et al., 2025) (Raffard et al. 2025). To account for heteroscedasticity (increasing residual variance with test temperature) in the metabolic rate data, we specified a variance power function (varPower) on test temperature (J. Pinheiro et al., 2025; J. Pinheiro & Bates, 2000).

In all models, we included all biologically relevant two- and three-way interactions between test temperature, temperature conditions (evolutionary conditions), and predation regime (evolutionary conditions), to test how the evolutionary conditions influenced trait variability interactively and whether these effects depended on the test temperature. Model significance was evaluated using likelihood ratio tests (LRTs) comparing nested models. We used backward selection by sequentially removing non-significant interactions (retaining lower-order terms when part of significant higher-order interactions). Confidence intervals were obtained by parametric bootstrapping with 50,000 simulations.

#### b. Evolutionary mechanisms underlying trait variability: F_ST_/P_ST_ analysis

##### Estimation of F_ST_

As an estimate of genetic drift, F_ST_ among experimental populations was estimated using an Analysis of Molecular Variance (AMOVA, Excoffier et al., 1992) that decomposes global genetic differentiation between and within populations. The ratio of the between variance component to the total molecular variance provides an estimate (expressed as a percentage) of the genetic variance explained among experimental populations.

##### Estimation of P_ST_

For each trait and experimental population, we estimated the mean and, when relevant, a parameter describing its response to test temperature. For body mass, only the mean for each experimental population was calculated, as this trait has not been measured at different test temperatures. For metabolic rate, we calculated, for each experimental population, the mean metabolic rate, and from the relationship between metabolic rate and test temperatures: the temperature of maximal metabolic rate (hereafter “temperature-dependent optimum metabolic rate”) and the thermal breadth across the experimental gradient (hereafter “thermal breadth of metabolic rate”). For decomposition, we estimated for each experimental population the mean decomposition rate and the slope of its relationship with test temperatures (hereafter “temperature-dependent decomposition rate”).

To assess trait differentiation (P_ST_) among the four evolutionary conditions (ambient/predation, ambient/predator-free, warm/predation, warm/predator-free), we set a linear mixed model independently for each variable cited above (mean body mass, mean metabolic rate, temperature-dependent optimum metabolic rate, thermal breadth of metabolic rate, mean decomposition rate and temperature-dependent decomposition rate). This model was an intercept-only model in which the random term was the experimental population identity nested within each evolutionary condition to which it belongs (populations nested within conditions). We extracted the estimate of the random term and assessed P_ST_ as the ratio (expressed as a percentage) between the among condition variance estimate to the total variance of a given trait. For P_ST_ values, we estimated 95% confidence intervals using a Jackknife procedure, as bootstrap resampling could fail to capture all evolutionary conditions. For F_ST_ values, we estimated 95% confidence intervals using bootstrap procedures on loci, where bootstrap resampling does not suffer from the same limitations as for P_ST_ (1000 bootstraps). We considered that P_ST_ and F_ST_ values were different from one another when their 95% CI did not overlap. We interpreted phenotypic divergence relative to neutral expectations: traits with P_ST_ ≈ F_ST_ were primarily influenced by genetic drift, P_ST_ < F_ST_ indicated stabilizing selection limiting divergence, and P_ST_ > F_ST_ reflected divergent selection among populations or plasticity.

## Results

### Trait variability: metabolic rate, body mass and decomposition rate

#### Metabolic rate

Metabolic rate varied with the test temperature (*p-value* < 0.0001, 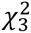 = 307.37; Figure 2), but was similar whatever the evolutionary conditions under which the experimental populations evolved (Table 1. Table S1). Regardless of the evolutionary condition under which populations evolved (temperature and predation regime), metabolic rate was the lowest at ∼13.33°C (metabolic rate predicted = 0.0097 mg O_2_ L^-^¹ min^-^¹, 95% CI [0.0045; 0.021], for an average-sized individual) and the highest at ∼27.22°C (metabolic rate predicted = 0.017 mg O_2_ L^-^¹ min^-^¹, 95% CI [0.0082; 0.038], for an average-sized individual).

**Figure 2.**
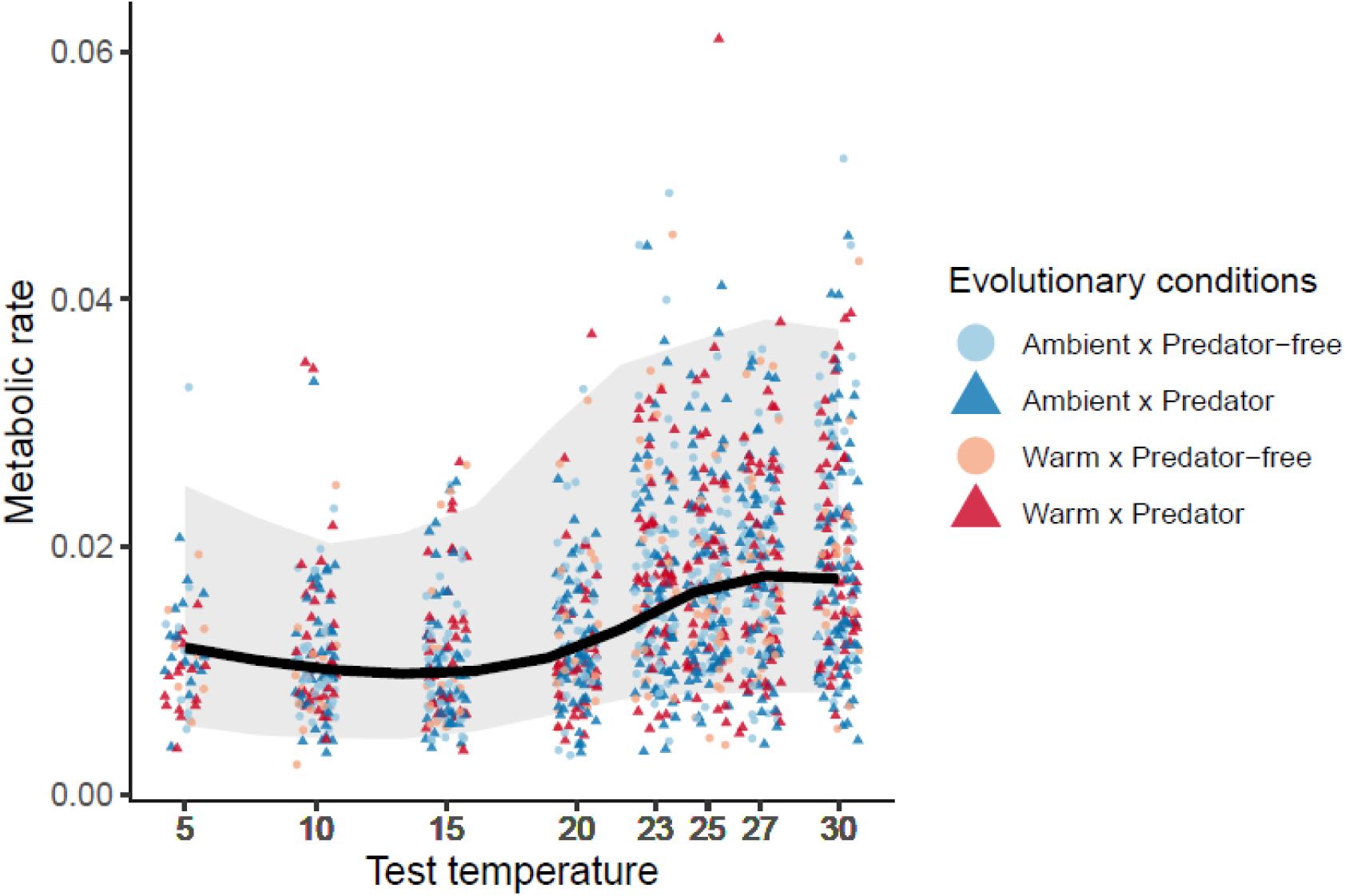
Metabolic rate depending on test temperature (x-axis: mg 0_2_.L^-^¹.min^-^¹; y-axis: °C). The metabolic rate corresponds to the slope of oxygen consumption (mg 0_2_.L^-^¹) over time (min). The solid line is the estimated mean relationship between mass-independent metabolic rate and test temperature from the best-fit linear mixed-effects model, with body mass held at its mean value. The shaded area is the 95% confidence interval of this estimate. Each point represents the metabolic rate of an individual whose color corresponds to the evolutionary condition. Model prediction (line) and raw data (points) were back-transformed from the log scale to the original scale for visualization.

**Table 1.** Likelihood ratio test (LRT) results for the effects of temperature condition, predation regime, and test temperature on metabolic rate, body mass, and decomposition rate. All models were linear mixed-effects models including experimental population as a random effect. While temperature condition and predation regime correspond to evolutionary conditions, test temperature corresponds to the experimental test temperature. For metabolic rate (log of the slope of oxygen consumption over time), log-transformed individual mass was included as a covariate, a variance structure was used to account for heteroscedasticity, and test temperature was modeled as a natural spline with three degrees of freedom. Significant p-values are shown in bold. Based on a backward model selection approach, non-significant effects were sequentially removed using an LRT test.

| Response | Effect | $\chi^2$ | df | *p*-value |
| --- | --- | --- | --- | --- |
| **Metabolic rate** |  |  |  |  |
|  | Temperature condition:Predation Regime:Test temperature | 0.940 | 3 | 0.816 |
|  | Temperature condition:Predation Regime | 2.970 | 1 | 0.085 |
|  | Predation Regime:Test temperature | 3.202 | 3 | 0.362 |
|  | Temperature condition:Test temperature | 3.012 | 3 | 0.390 |
|  | Temperature condition | 0.791 | 1 | 0.374 |
|  | Predation regime | 1.033 | 1 | 0.309 |
|  | **Test temperature** | **307.26** | **3** | **<.0001** |
| **Body Mass** |  |  |  |  |
|  | Temperature condition:Predation Regime | 1.689 | 1 | 0.194 |
|  | Temperature condition | 0.456 | 1 | 0.500 |
|  | Predation regime | 0.774 | 1 | 0.379 |
| **Decomposition rate** |  |  |  |  |
|  | Temperature condition:Predation Regime:Test temperature | 3.732 | 1 | 0.053 |
|  | Temperature condition:Predation Regime | 0.319 | 1 | 0.572 |
|  | **Predation Regime:Test temperature** | **18.435** | **1** | **0.000** |
|  | **Temperature condition:Test temperature** | **7.003** | **1** | **0.008** |

#### Body mass

Body mass was not significantly different across evolutionary conditions under which the experimental populations evolved, and we did not identify any particular trend (Figure 3-A, Table 1, Table S2). Body size showed comparable trends across evolutionary conditions (Figure 3-B).

**Figure 3.**
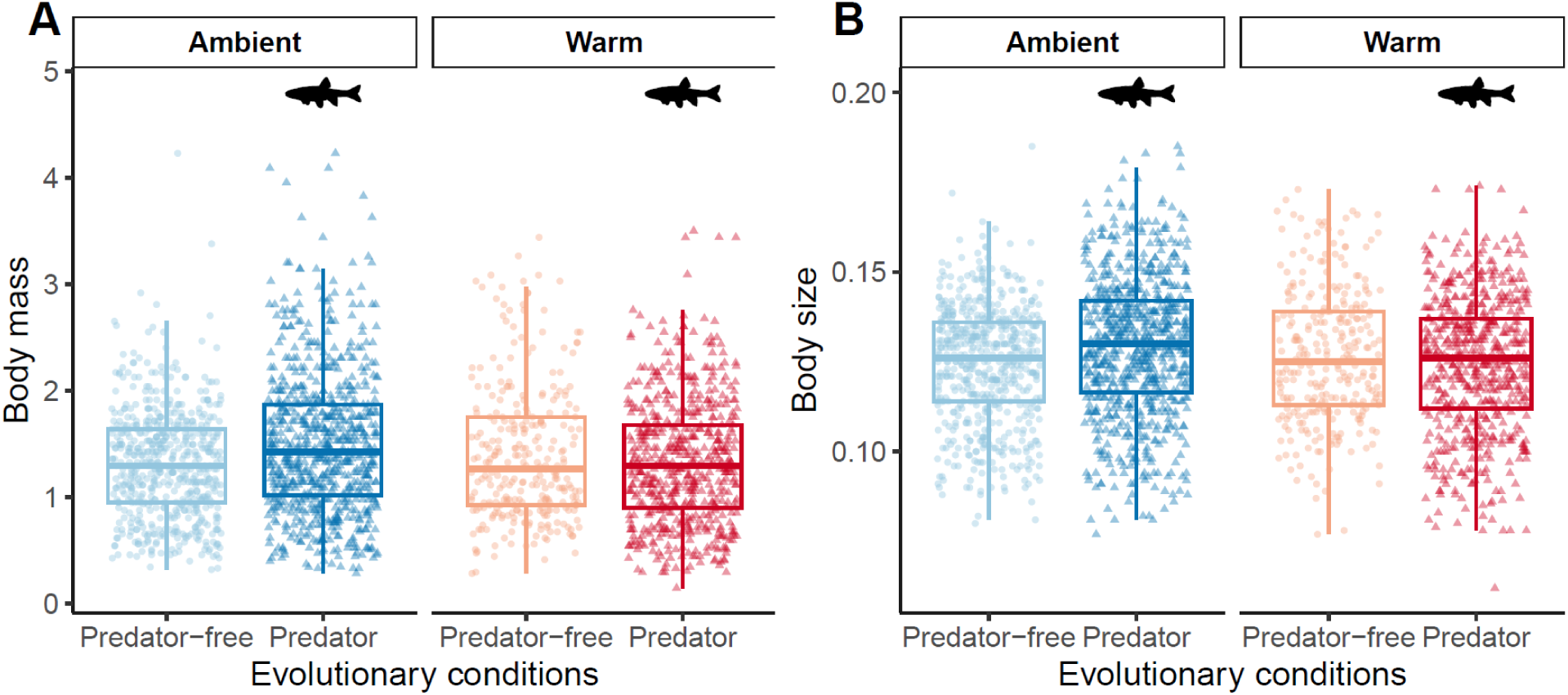
(A) Body mass (y-axis: mg) and (B) Body size of individuals (y-axis: cm) as a function of the conditions under which they evolved. Each point represents an individual whose color and shape correspond to the evolutionary conditions under which they evolved. Body mass was estimated for each individual using the body size-body mass relationship (see Figure S1).

#### Decomposition rate

Decomposition rate was significantly affected by the interaction of test temperature with both evolutionary temperature conditions (*p-value* = 0.008, 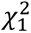 = 7.00, Table 1) and evolutionary predation regime (*p-value* < 0.0001, 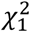 = 18.44, Table 1). Indeed, the decomposition rate increased with test temperature, but this increase varied significantly among experimental populations. The influence of the test temperature on decomposition rate increased when populations evolved without predation (increase in the slope of 2.3e-04 with se = 5.3e-05) and increased when populations evolved in warm conditions (increase in the slope of 1.4e-04 with se = 5.4e-05, Table S3). Particularly, the increase in decomposition rate with test temperature was more marked for populations having evolved in warm and predator-free conditions (Figure 4, with a slope of 9.5e-4 with se=5.3e-05). At 27°C, where the effects of evolutionary conditions were most pronounced, the decomposition rate of warm x predator-free populations was predicted at 0.032, while that of ambient x predator populations was predicted at 0.026.

**Figure 4.**
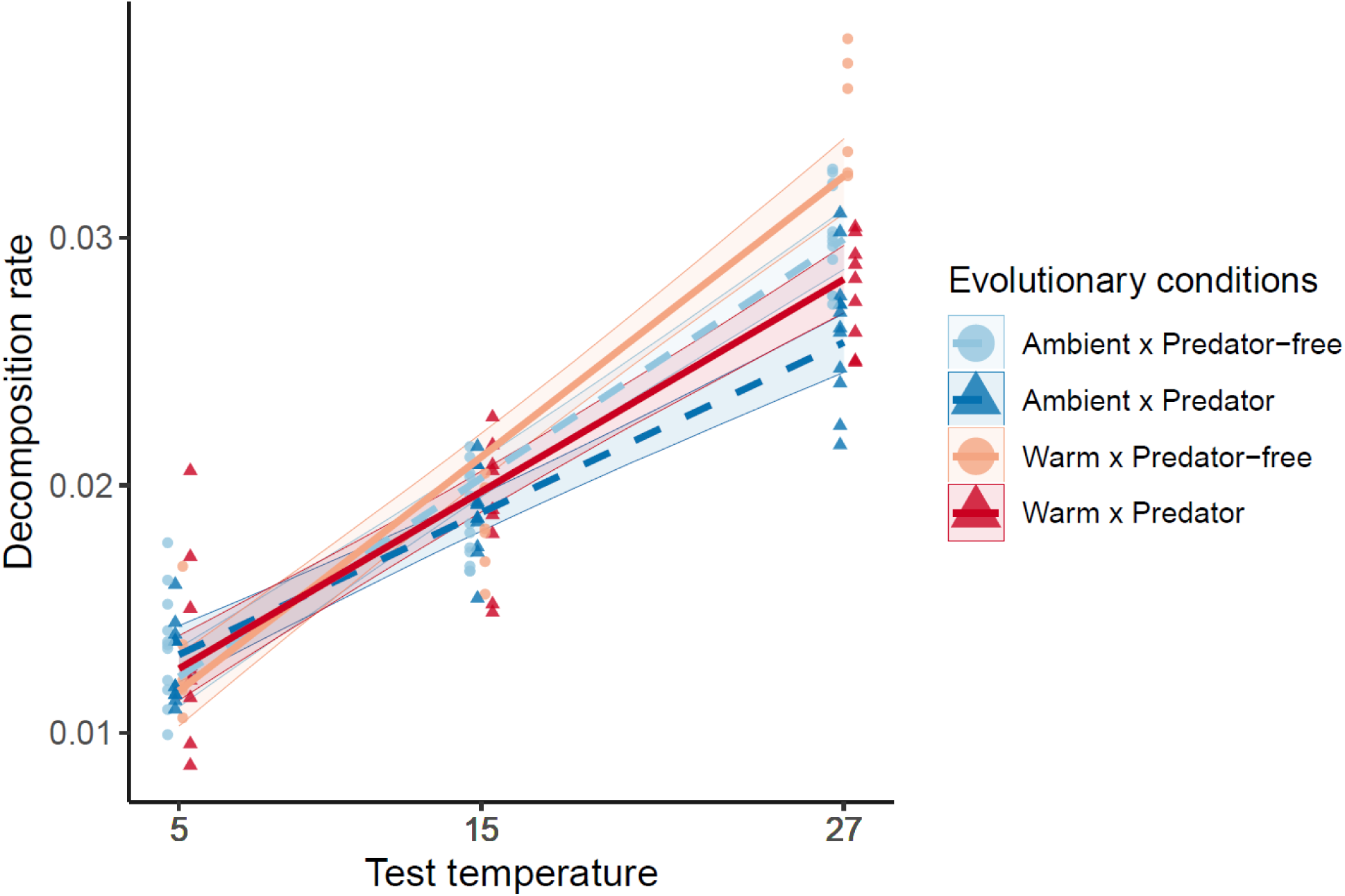
Decomposition rate depending on test temperature and evolutionary conditions (y-axis: day^-^¹; x-axis: °C). The points represent raw data for each evolutionary condition and are slightly staggered on the temperature axis to reduce overlap (test temperature is either 5°C, 15°C or 27°C). The solid lines are the estimated mean relationship from the best-fit linear mixed-effects model. The shaded area is the 95% confidence interval of this estimate. The experimental populations evolved either in warm temperature (red with a solid line) or ambient temperature (blue with a dotted line), and either with the presence of a predator (dark colors with triangular points) or without a predator (light colors with circular points).

### Evolutionary mechanisms underlying trait variability: FST/PST analysis

Based on the F_ST_ estimation, we assumed that the amount of trait variability imputable to genetic drift would be 21.67% (95% CI: 20.98-22.34). With several P_ST_ values differing significantly from this neutral baseline, we showed that phenotypic differentiation could not be explained by genetic drift only (Figure 5). Indeed, the P_ST_ values for mean body mass, temperature-dependent optimum metabolic rate, mean decomposition rate and temperature-dependent decomposition rate were significantly different from the neutral baseline. More specifically, the P_ST_ associated with mean body mass and temperature-dependent optimum metabolic rate was significantly lower than the F_ST_ value: this suggests that for these two traits stabilizing selection likely constrained phenotypic divergence among populations, potentially driven by strong selective pressures maintaining optimal body size and metabolic performance under different conditions. On the contrary, the P_ST_ associated with decomposition rate (mean decomposition rate and temperature-dependent decomposition rate) were significantly higher than the F_ST_ value with a very strong divergence, indicating that for these two traits, divergent selection or local adaptation promoted phenotypic differentiation among populations. For the mean metabolic rate and thermal breadth of metabolic rate, P_ST_ statistically equaled F_ST_, indicating that drift was the major driver of phenotypic trait divergence.

**Figure 5.**
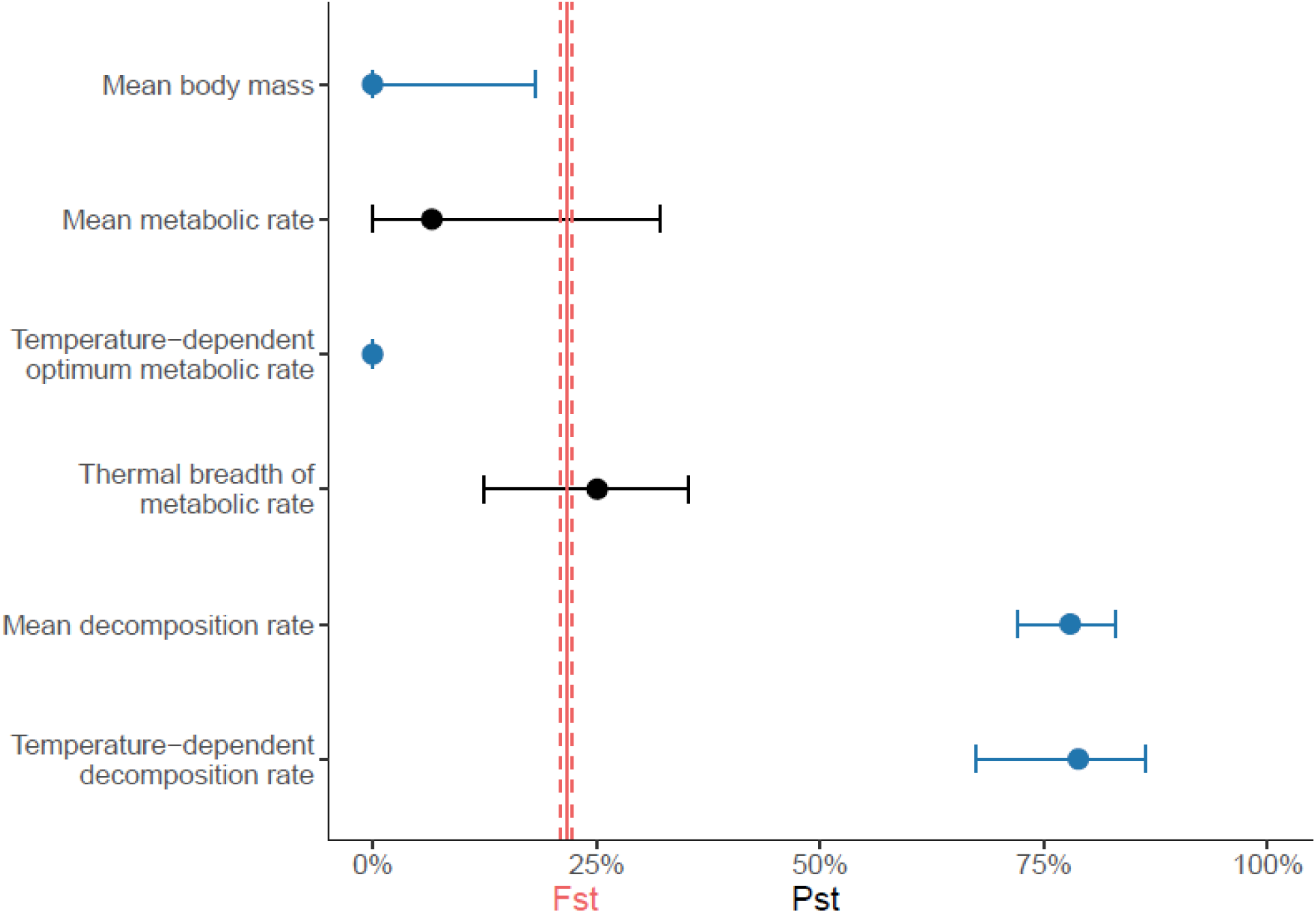
Comparison of genetic and trait differentiation values. The 95% confidence interval of the F_ST_ is represented by the red dotted vertical lines. The 95% confidence interval of the P_ST_ is represented by the horizontal lines. When the confidence intervals of the P_ST_ and F_ST_ do not overlap, P_ST_ is considered significantly different from the F_ST_ and its confidence intervals are represented in blue. For mean body mass and temperature-dependent optimum metabolic rate, error bars are not visible because their confidence intervals are smaller than the plot symbols.

## Discussion

Here, we evaluated how the evolutionary conditions in which *Asellus aquaticus* populations evolve shape both functional traits and the ecosystem function. We found that the evolutionary conditions (temperature and predation regime) did not affect metabolic rate nor body mass, both functional traits that are thought to influence ecosystem functioning. In contrast, evolutionary conditions influenced decomposition rate, which is the function of the ecosystem itself.

Furthermore, both metabolic rate and decomposition rate varied along a temperature gradient (i.e., the test temperature). We also showed that neutral processes mainly contributed to variation in metabolic rate, whereas adaptive processes shaped variation in body mass and decomposition rate but potentially due to opposite reasons.

Our results indicate that evolutionary conditions can modify the ecosystem functioning in a few generations (rapid evolution), without necessarily altering functional traits. Indeed, when *Asellus* populations evolve in warm conditions, they evolve toward a higher ability to decompose dead leaves, which seems to be sustained by natural selection and/or phenotypic plasticity (based on the importance of selection revealed across all evolutionary conditions by P_ST_ and F_ST_ analyses). Because a common garden design was used between the mesocosm experimental evolution and the measurements minimizing environmental plastic effects, the differences observed among evolutionary conditions likely reflect adaptive responses, either through genetic changes or genotype-by-environment interactions. In populations evolving in warm temperature, individuals with a greater ability to decompose at high temperatures would have been selected during the mesocosm experiment. As expected, selection for a highest ability to decompose dead leaves in the warm climate could be explained by an increase in energetic needs in these populations (Martínez et al., 2014). Also, in line with our initial hypotheses, populations that had evolved under predation pressure, either in warm or ambient conditions, systematically had a reduced ability to decompose organic matter (in particular when measured in warm temperature), likely reflecting behavioral adjustments such as reduced feeding activity to minimize predation risk (Raffard et al., 2017). Results from the P_st_/F_st_ analysis confirm that decomposition rate quickly responded to directional selection for each of the evolutionary conditions. Decomposition rate in macro-decomposers depends on multiple traits, such as physiological, metabolic, and behavioral traits. Behavioral traits, such as food choices or time spent feeding, can evolve much faster than other traits such as metabolic rate, especially in the presence of predators (Hawlena & Schmitz, 2010), allowing fast adaptive responses.

Given that the decomposition of organic matter is the main source of energy for most ecosystems such as rivers or wooded lakes, this result highlights the influence that evolutionary conditions can have on a function that will have a strong impact on ecosystem dynamics. This example of eco-evolutionary interaction was also highlighted by Dossena et al. (2012), where the evolution of benthic communities in aquatic mesocosms showed altered decomposition rates and community structure in treatments exposed to warm temperatures.

Contrary to our expectations, neither metabolic rate nor body mass evolved in response to warming or predation. Regarding body mass, the lack of evolution contrasts with the reduction in size expected in ectotherms in response to warming, as predicted by the Temperature–Size Rule (Daufresne et al., 2009; Heneghan et al., 2019; Wootton et al., 2022). Our results indicate that variability of body mass in our experiment was not solely explained by drift, but also largely by selection. Body mass is an extremely constrained and conserved trait, potentially under stabilizing selection as suggested here. In crustaceans, moulting and therefore size are under very strong genetic control, limiting a rapid response to environmental pressures. Rapid changes are hampered by the need to modify moulting regulators, which can be costly or pleiotropic (Campli et al., 2024). In the context of a highly constrained trait, evolutionary shifts in body mass may take longer than the 6–8 generations of *Asellus* in this experiment. Consistent with this, similar experiments in *Daphnia* detected changes in body mass in response to temperature or the presence of a predator only after several tens of generations (Cavalheri et al., 2019; Doorslaer et al., 2010; Wang et al., 2023).

Regarding metabolic rate, the absence of evolutionary change is surprising, as local thermal adaptation is expected to shape metabolic rates according to the Metabolic Cold Adaptation theory, among other hypotheses (Angilletta, 2009; Lardies et al., 2004; Somero, 2012; Williams et al., 2016). However, similar patterns have been reported in other long-term experimental evolution studies (Alberto-Payet et al., 2022; Andreassen et al., 2025). For instance, after nine years of experimental evolution, populations of the medaka fish *Oryzias latipes* evolving over a dozen generations under warm and cold temperatures did not show differences in resting metabolic rate (Alberto-Payet et al., 2022). This surprising lack of evolution could be explained by the specific trait measured in our experiment: the routine metabolic rate. In Zebrafish, populations selected for thermal tolerance over multiple generations show improved performance under warming, such as cooling tolerance, without altering routine metabolic rate (Andreassen et al., 2025). Additional experiments measuring a panel of traits such as individual thermal tolerance, could help to draw conclusions on this point. However, the lack of evolution in such “classical ecological traits” raises questions about whether such traits reliably capture the organismal determinants of ecosystem functioning.

Together, these results reveal a disconnect between functional traits assumed to underlie ecosystem functions and the ecosystem function. While body size and metabolism are widely used as ecological proxies, they showed weak rapid evolutionary responses, whereas the ecosystem function itself—decomposition rate—evolved rapidly. This demonstrates that focusing solely on functional traits can obscure key eco-evolutionary dynamics, and highlights the value of directly measuring ecosystem functions rather than relying on trait-based assumptions. This conclusion aligns with recent critiques of trait-based ecology, which argue that many commonly used functional traits lack demonstrated mechanistic links to ecosystem processes and that a function-first approach is needed to identify traits truly relevant to ecosystem functioning (Streit & Bellwood, 2023).

Both metabolic rate and decomposition rate showed phenotypic plasticity, as measured values varied along the study temperature gradient. Indeed, the metabolic rate varied non-linearly with temperature at which it was measured, with high rates observed at stressful temperatures (below 10°C or above 25°C). Such patterns are consistent with other studies of thermal stress in aquatic organisms, where both the metabolic and physiological responses exhibit clear thresholds and shifts, often becoming nonlinear at extreme temperatures (Pörtner et al. 2017). The decomposition rate increased with the temperature at which it was measured, a trend observed in several aquatic ecosystems where higher temperatures accelerate both microbial and detritivore-mediated decomposition rates (Bernabé et al., 2018). This variation was even higher depending on the evolutionary conditions, highlighting the importance of local adaptation in shaping these thermal responses. As demonstrated in our experiment, populations from warmer environments often exhibit higher phenotypic plasticity, potentially allowing for more robust responses to temperature variability (Yampolsky et al., 2014). Similar evolutionary shifts toward broader reaction norms have been reported in other systems, suggesting that environmental variability can select for increased phenotypic plasticity, as shown for instance in marine diatoms evolving under warm conditions (Schaum et al., 2018). More broadly, the evolution of plasticity is increasingly recognized as a rapid adaptive response to climate change, allowing populations to cope with novel or variable environments (Kelly, 2019).

Global change, including both warming and other stressors, is expected to amplify these temperature-dependent shifts, altering biogeochemical cycles and food web dynamics across ecosystems (Luo et al., 2025; Pan et al., 2024). Integrating thermal reaction norms and plasticity into predictive models is therefore essential to forecast how aquatic ecosystem functioning will respond to ongoing climate warming (Kordas et al., 2022).

Our study is limited by two factors related to our experimental design. First, the relatively short duration of the mesocosm experiment constrained the number of generations (6–8) over which evolutionary change could occur. Although this timescale was sufficient to detect evolution in decomposition rate, a longer experiment would have increased the likelihood of observing responses in traits with weaker selection gradients, more complex genetic bases or strong genetic constraints such as the body mass (Kawecki et al., 2012). Second, the common-garden phase, which lasts 48 hours or 6 weeks depending on the trait, may not have fully eliminated environmentally induced effects (Fry, 2003). Although this approach aligns with established *Asellus* protocols (Lürig et al., 2019), some residual plastic or transgenerational effects may persist after only partial rearing (Kawecki & Ebert, 2004). In summary, increasing the duration of the common-garden phase would have further reduced residual plastic effects.

However, because the total duration of the project was limited, extending the common-garden phase would have reduced the duration of the mesocosm experiment, and consequently, the number of generations available for evolutionary change. Our design therefore represents a trade-off between controlling non-genetic effects and allowing sufficient time for evolutionary responses to occur.

To conclude, our findings indicate how global change–related pressures can foster rapid evolutionary changes in a key functional trait, namely the ability of macro-organisms to decompose dead organic matter. Given the ecological importance of this trait, our study suggests that these evolutionary changes could have cascading effects on the entire food web. We show that measuring directly the ecosystem function is critical for predicting these ecosystem dynamics. Indeed, despite the short time scales, our study suggests that functions such as the decomposition rate can evolve upon evolutionary mechanisms or plasticity and alter ecosystem functioning, even when traits commonly used as functional proxies remain stable throughout the same period. Integrating such dynamics into predictive models would be essential for forecasting ecosystem responses under global change.

## Supporting information

SupMat

## Notes

### Competing Interest Statement

The authors have declared no competing interest.

### Summary of Updates

Add one author in the list of the author

