## Supplementary material for "Warming and predation drive rapid evolution of ecosystem functioning but not functional traits": SupMat

### Supplements

***Table S1 Summary of the best-fitting linear mixed-effects model describing variation in metabolic rate.*** *The best-fit model was obtained following stepwise model selection using likelihood ratio tests (LRT) to remove non-significant terms. The model included test temperature, modeled as a natural spline with three degrees of freedom (ns(TempExpNum, 3)), and log-transformed individual mass (log(Mass_Ind)) as a covariate. Population (Pop) was included as a random intercept, and a variance structure (varPower) was applied to account for heteroscedasticity as a function of test temperature. The model was specified as: lme(log(Slope_Pos) ~ ns(TempExpNum, 3) + log(Mass_Ind), random = ~1 | Pop, weights = varPower(form = ~ TempExpNum)).*

| **Fixed effects** | **Estimate** | **Std. Error** | **t-value** |
|---|---|---|---|
| (Intercept) | -4.548184 | 0.04999703 | -90.96908 |
| ns(TempExpNum,3)1 | 0.552541 | 0.04311947 | 12.81419 |
| ns(TempExpNum,3)2 | 0.055413 | 0.11701444 | 0.47355 |
| ns(TempExpNum,3)3 | 0.612546 | 0.03689561 | 16.60213 |
| log(Mass_Ind) | 0.278766 | 0.03022373 | 9.22343 |

***Table S2 Summary of the best-fitting linear mixed-effects model describing variation in individual body mass.*** *The best-fit model was obtained following stepwise model selection using likelihood ratio tests (LRT) to remove non-significant terms. The final model included only an intercept and a random effect of population (*Pop*), accounting for among-population variability in mean body mass. The model was specified as: lmer(Mass_Ind ~ 1 + (1 | Pop)).*

| **Fixed effects** | **Estimate** | **Std. Error** | **t-value** |
|---|---|---|---|
| (Intercept) | 1.39E+00 | 5.26E-02 | 26.44 |

***Table S3 Summary of the best-fitting linear mixed-effects model explaining decomposition rate.*** *The best-fit model was obtained following stepwise model selection using likelihood ratio tests (LRT) to remove non-significant terms. The selected model included test temperature (*TempExpNum*), predation regime (*Predpop*), and temperature condition (*Tpop*), as well as their two-way interactions with* TempExpNum*. Population (*Pop*) was included as a random effect, though its estimated variance was null. The model was specified as: lmer(Deg_Rate_Raffard ~ TempExpNum:Predpop + TempExpNum:Tpop + TempExpNum + Tpop + Predpop + (1 | Pop)).*

| **Fixed effects** | **Estimate** | **Std. Error** | **t-value** |
|---|---|---|---|
| (Intercept) | 8.25E-03 | 7.88E-04 | 10.462 |
| TempExpNum | 8.03E-04 | 4.32E-05 | 18.607 |
| TpopWarm | -1.28E-03 | 9.80E-04 | -1.302 |
| PredpopPred | 2.05E-03 | 9.62E-04 | 2.132 |
| TempExpNum:PredpopPred | -2.30E-04 | 5.30E-05 | -4.342 |
| TempExpNum:TpopWarm | 1.42E-04 | 5.41E-05 | 2.616 |

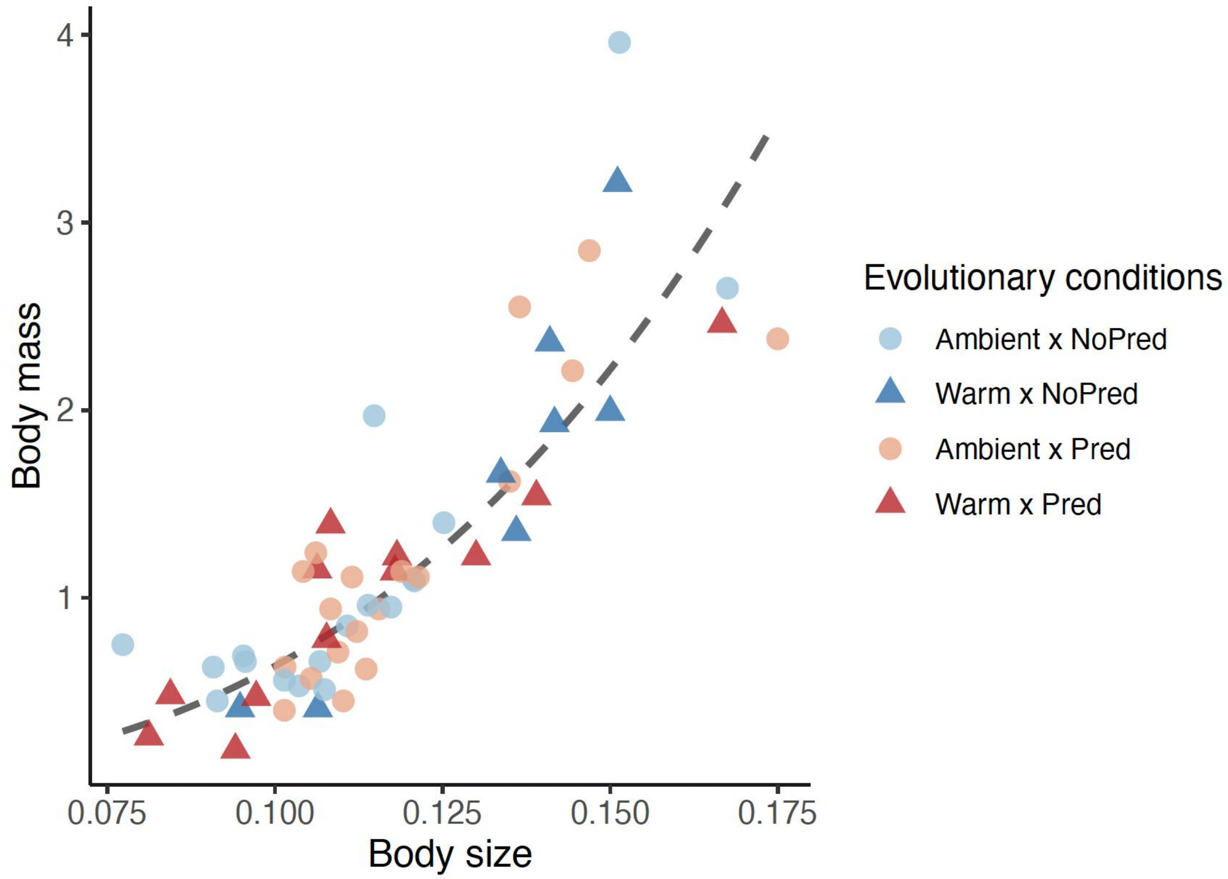


***Figure S1. Relation between body mass and body size.*** *Each point represents an individual, with color and shape indicating the conditions under which they evolved. The dotted line shows the fitted allometric relationship (body mass = 765.99 × body size$^{3.08}$) derived from the log–log model log(body mass) = 6.64 + 3.08 × log(body size) ; with an $R^2$ = 0.721. The body mass of the individuals presented in this study (Figure 2) was estimated using this relationship estimated for these individuals (body mass = 765.99 × body size$^{3.08}$).*

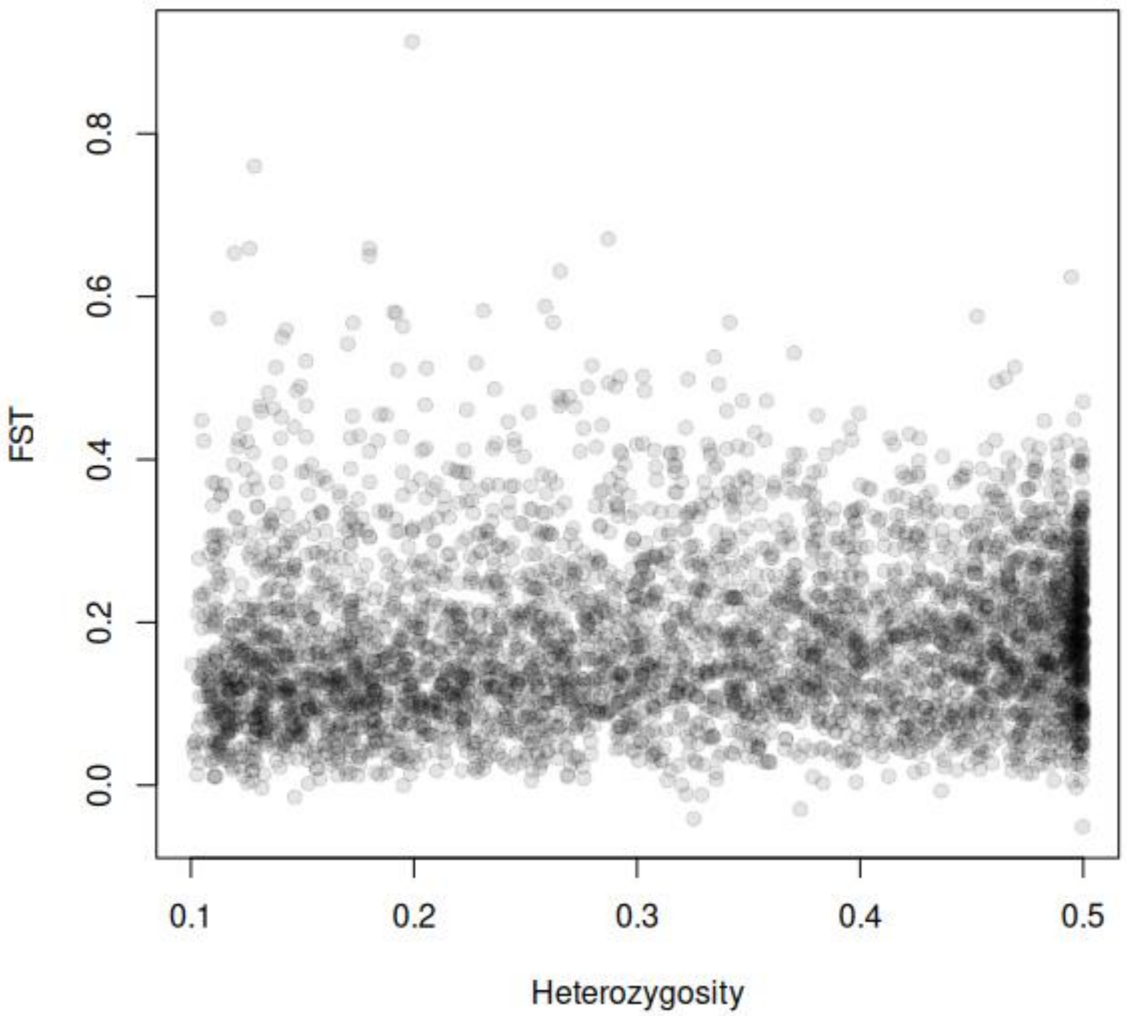


**Figure S2 *Relation between Fst and Heterozygosity.*** *Each point represents a locus.* The OutFLANK approach *uses likelihood on a trimmed distribution of $F_{ST}$ values to infer the distribution of $F_{ST}$ for neutral markers (Whitlock and Lotterhos 2015). This distribution is then used to assign q-values to each locus to detect outliers that may be under selection.* Based on this approach, no $F_{ST}$ outliers were detected.
